# A proposed chaperone of the bacterial type VI secretion system functions to constrain a self-identity protein

**DOI:** 10.1101/213785

**Authors:** Martha A. Zepeda-Rivera, Christina C. Saak, Karine A. Gibbs

## Abstract

The opportunistic bacterial uropathogen *Proteus mirabilis* can communicate identity through the export of the self-identity protein, IdsD, via the type VI secretion (T6S) system. Expression of the *ids* genes provides a fitness advantage during polymicrobial infections in a mouse infection model. Here we provide an answer to the unresolved question of how the activity of a T6S substrate, such as IdsD, is regulated before export. We demonstrate that IdsD is found in clusters that form independently of the T6S machinery and activity. We show that the protein IdsC, which is a member of the proposed DUF4123 chaperone family, is essential for the stability of these clusters as well as the IdsD protein itself. And we provide evidence that amino acid disruptions in IdsC are sufficient to disrupt IdsD export but not IdsD localization into stable subcellular clusters, strongly supporting that IdsC functions in at least two different ways: IdsD stabilization and IdsD export. We propose that IdsC, and likely other DUF4123-containing proteins, function to regulate T6S substrates before export by both stabilizing the protein and mediating export at the T6S machinery.

## Introduction

Recognition of self versus non-self is a broadly observed behavior. For single-celled organisms, recognition can allow organisms of close genetic relatedness to cooperate to access benefits and behaviors that individual cells could otherwise not achieve. There are many described mechanisms for these recognition behaviors, such as the exchanges of lethal proteins (1–8), of freely diffusible molecules (9–14), and of non-lethal identity proteins (15, 16). For several non-lethal mechanisms, such as in the amoeba *Dictyostelium discoideum* during spore formation (17–19) and in the bacterium *Myxococcus xanthus* during outer membrane exchange (20–23), communicating identity among sibling cells depends on cell-to-cell contact wherein binding interactions between surface-exposed proteins on neighboring cells signal the presence or absence of kin. Similarly, self recognition in the bacterium *Proteus mirabilis* depends on cell-to-cell contact while migrating on a surface, a behavior termed "swarming" (15). In contrast to *D. discoideum* and *M. xanthus*, identity in *P. mirabilis* is conveyed by the transfer of a self-identity protein, IdsD, into a neighboring cell where it interacts with its identity partner protein, IdsE (16, 24). Strain-specific variable regions, comprised of several amino acids, within IdsD and IdsE confer binding specificity; cognate IdsD and IdsE variants bind, while non-cognate IdsD and IdsE variants do not bind (24). Intriguingly, IdsD and IdsE are co-regulated and predicted to localize to the inner membrane of *P. mirabilis* cells (24, 25), yet only the binding status of IdsD-IdsE in the recipient cell contributes to self recognition. IdsD that is bound by IdsE allows proficient population swarming, while IdsD that is not bound results in restricted population swarming (16). These findings indicate that IdsD and IdsE likely do not interact in the producing (donor) cell before export and provoked the question of how IdsD activity is regulated in the donor cell to prevent self-restriction.

IdsD export depends on a type VI secretion (T6S) system (26). Intriguingly, expression of both *ids* and *t6s* genes provides a fitness advantage for *P. mirabilis* during polymicrobial infections in a mouse infection model (27). T6S systems are multi-protein, cell-envelope spanning transport machineries found broadly among gram-negative bacteria. These T6S systems generally act as conduits through which substrates are sent from the interior of a donor cell into the interior of a recipient cell (26, 28–33). For export out of the donor cell, T6S substrates often interact with components of the T6S transport machinery as well as associated proteins, including the proposed DUF4123-containing protein chaperones as well as VgrG- and PAAR-containing proteins (34–41). Once in the recipient cell, many of the T6S substrates, termed “effectors”, have binding partners ("immunity proteins") that neutralize the effector's activity if the donor and recipient cells are related (30, 33, 42, 43). For many of these cognate two-partner proteins, the immunity protein is predicted to also prevent effector activity in donor cells. IdsD’s interaction partners for export out of a donor cell are yet unknown. Given this, and that IdsD does not interact with IdsE before transport, a pressing question is how IdsD activity is regulated in donor cells before export.

Here we have combined biochemical, genetic, and imaging techniques to address whether protein-protein interactions regulate IdsD before export. We demonstrate that a third protein, IdsC, which contains a predicted DUF4123 domain, is essential for the stabilization of IdsD into subcellular clusters in the donor cell independently of export via T6S. Formation and localization of IdsD-containing clusters was unaffected by the absence of IdsE. We further show that strain-specific single amino acid variations across IdsC do not impact the IdsC-IdsD binding interaction or formation of IdsD clusters, but do disrupt IdsD export. Taken together, these data support that there are minimally two chaperone functions for IdsC: IdsD stabilization and IdsD export. This predicted chaperone activity provides one explanation for why IdsD and IdsE do not bind in a single cell and challenges the current perception that the substrate-specific immunity protein suppresses T6S substrate activity in donor cells.

## Results

### IdsD subcellular localization is independent of export

The subcellular location of IdsD, as well as other T6S substrates in *P. mirabilis*, was previously unknown; however, we predicted that IdsD would localize with the T6S machinery for transport. To resolve the subcellular localization of IdsD, we engineered a functional fusion of mKate2 to IdsD (mKate-IdsD) that replaced unlabeled IdsD in *P. mirabilis* producing a functional N-terminal FLAG epitope-tagged IdsC (Figure SF1). Fluorescence associated with mKate-IdsD was found as discrete foci, often near the poles, in a subset of cells (Figure 1A). IdsD localization with respect to the T6S machinery was analyzed by introducing a chromosomal fusion of superfolder Green Fluorescent Protein (sfGFP) to a sheath-encoding gene, *tssB* (44). Briefly, T6S requires the contraction of a subcellular sheath comprised of TssB and TssC that plunges an interior tube of Hcp protein hexamers across the cell envelope through a core membrane complex consisting of several proteins including TssM (29, 36, 37, 45–49). The baseplate protein, TssK, links sheath assembly and the core membrane complex (50, 51). In swarming cells producing mKate-IdsD and TssB-sfGFP, 26% of the mKate-IdsD associated foci were found proximal to the fluorescence associated with TssB-sfGFP. However, the other 74% of IdsD-associated foci were not found proximal to any of the T6S machineries within a given cell (Figure SF2), raising the possibility that these IdsD foci formed independently of the T6S machinery or active transport.

**Figure 1.**
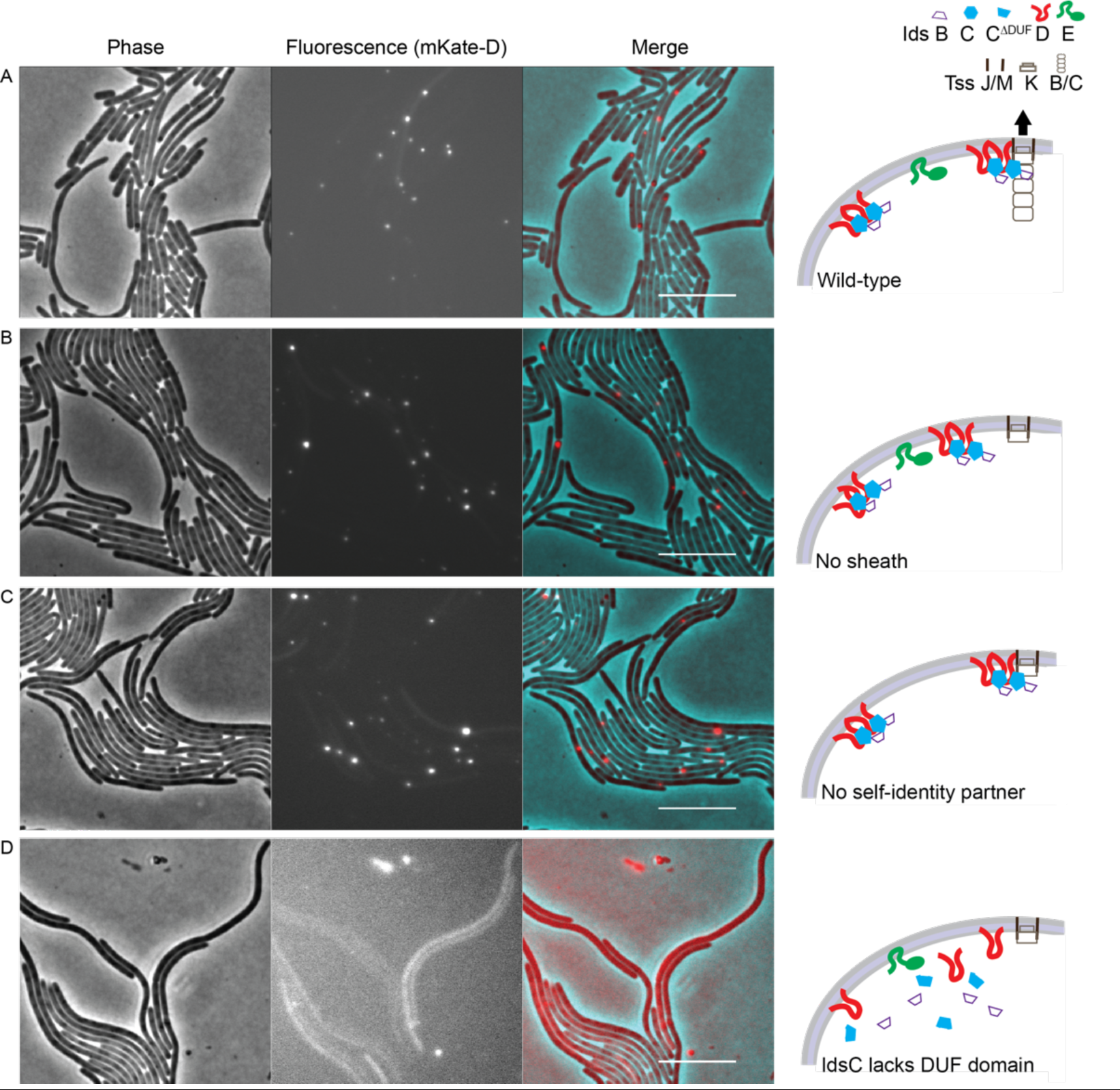
IdsD localization into subcellular clusters is dependent on IdsC but not IdsE (partner identity protein). A *P. mirabilis* strain in which FLAG-IdsC and mKate-IdsD replace the native IdsC and IdsD; all of the other Ids and T6S proteins are produced unless otherwise noted. Left, Phase. Middle, fluorescence in the RFP channel for mKate2. Right, false-colored overlay in which for contrast, mKate-IdsD fluorescence is in red, and phase is in cyan. All scale bars are 10 µm. Illustrations to the right of images depict models for IdsD localization at the membrane. Three main components of the T6S (the baseplate, the membrane-complex, and sheath) are shown. Black arrow represents tested IdsA (Hcp homolog) secretion, which is a hallmark of T6S activity (16, 26). (A) *P. mirabilis* strains producing FLAG-IdsC and mKate-IdsD. Fluorescence associated mKate-IdsD foci overlap at a 26% frequency with fluorescence associated with the T6S sheath (Figure SF2). (B) A *P. mirabilis* strain producing FLAG-IdsC and mKate-IdsD in which T6S sheath formation is disrupted by a mutation in a TssB homolog (16). (C-D) A *P. mirabilis* strain with disrupted sheath formation and in which either (C) the partner identity protein, IdsE, is deleted as previously described (16), or (D) the DUF4123 domain of IdsC is removed.

To address this hypothesis, we determined the localization of IdsD in donor cells, using strains containing distinct disruptions of the singular T6S machinery. Sheath formation was abrogated using a single point mutation into the gene encoding the TssB homolog (16, 44). The core membrane complex was impaired by disrupting the gene encoding the TssM homolog (26). Baseplate formation was disrupted using two independent point mutations into the gene encoding the TssK homolog. One mutation (TssK_null_) prevented sheath formation (Figure SF3) and secretion (Figure SF4). The other mutation (TssK_partial_) resulted in reduced sheath formation (Figure SF3) and secretion of Hcp homologs but not of IdsD (Figure SF4). The four mutant strains impaired sheath formation and IdsD secretion to different extents (Figures SF3, SF4) (16, 26, 44). In all of these populations, mKate-IdsD-associated fluorescence was patterned as foci along the cell periphery (Figure1B, SF3). Therefore, neither the T6S machinery nor active transport was necessary for the subcellular localization of IdsD.

### IdsD subcellular localization depends on IdsC (DUF4123) but not IdsE

Two additional proteins required for T6S substrate export are VgrG and DUF4123-containing proteins. VgrG proteins are themselves often decorated by PAAR-proteins, cap the Hcp tube, interact with the TssK protein, and guide substrate export (34, 35, 37, 38). DUF4123-containing proteins, which also bind VgrG proteins and T6S substrates, are predicted chaperones of T6S systems (35, 41). Within the *ids* operon, *idsC* encodes for a DUF4123-containing protein that is required for IdsD self-recognition activity (15, 25, 26). We predicted that IdsC might be essential for the subcellular localization of IdsD. Furthermore, we wanted to examine whether IdsE, the identity protein interaction partner of IdsD, was necessary for IdsD subcellular localization.

Given that none of the four T6S-deficient strains impacted IdsD localization, we used the TssB mutant strain as a representative donor-only population to test IdsD localization in donor cells lacking IdsC or IdsE. In strains where *idsE* was deleted (16), IdsD-associated fluorescence was again observed as discrete foci along the cell (Figures 1C). IdsE was not necessary for the subcellular localization of IdsD.

We next tested whether IdsC was essential for the localization of IdsD. We disrupted FLAG-IdsC by removing the DUF4123 domain (Figure 3A), resulting in FLAG-IdsC^∆DUF^. The mKate-IdsD-associated fluorescence in cells producing FLAG-IdsC^∆DUF^ was diffuse across the cell interior with few to no foci visible (Figure 1D). To exclude the possibility that this dispersed fluorescence was due to cleavage of mKate2 from IdsD in the FLAG-IdsC^∆DUF^ strain, we performed western blot analysis on whole cell extracts from both the FLAG-IdsC^∆DUF^ and FLAG-IdsC strains producing mKate-IdsD. A band corresponding to the mKate-IdsD fusion was readily observed, while bands corresponding to mKate2 or IdsD alone were minimally found (Figure 2A). We concluded that cleavage of mKate-IdsD did not explain the dispersed fluorescence, and as such, full-length IdsC was necessary for the presence of mKate-IdsD foci.

**Figure 2.**
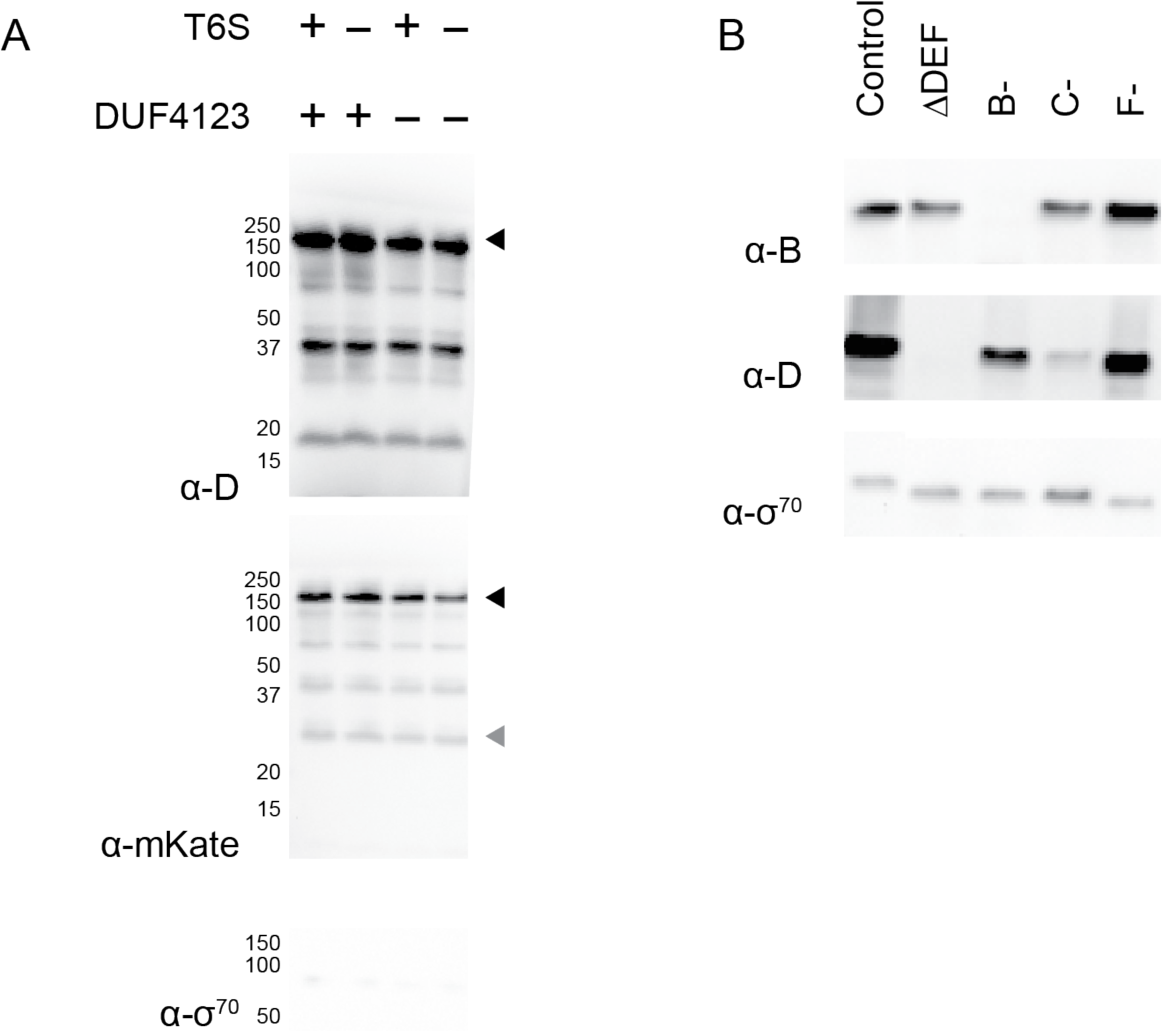
IdsC is essential for IdsD protein stability. (A) Whole cell lysates were collected from *P. mirabilis* strains with functional or disrupted T6S (16, 44) producing mKate-IdsD in conjunction with either FLAG-IdsC or FLAG-IdsC^ΔDUF^. Analysis was performed using polyclonal anti-IdsD, polyclonal anti-mKate, and monoclonal anti-sigma70 antibodies. Black arrow indicates size of mKate-IdsD fusion protein. Gray arrow indicates size of mKate protein. (B) Whole cell lysates were collected from *P. mirabilis* strains cells expressing differentially modified pIds_BB_ vectors (15), resulting in Ids protein deletions or disruptions. Lysates were normalized to OD_600_=1 and resuspended in loading buffer. 10 µl of each sample was run on a 12% tris-tricine gel and analyzed via western blot. Analysis was performed using polyclonal anti-IdsB, polyclonal anti-IdsD, and monoclonal anti-sigma70 antibodies. Labels above blot indicate samples; control is unmodified pIds_BB_.

**Figure 3.**
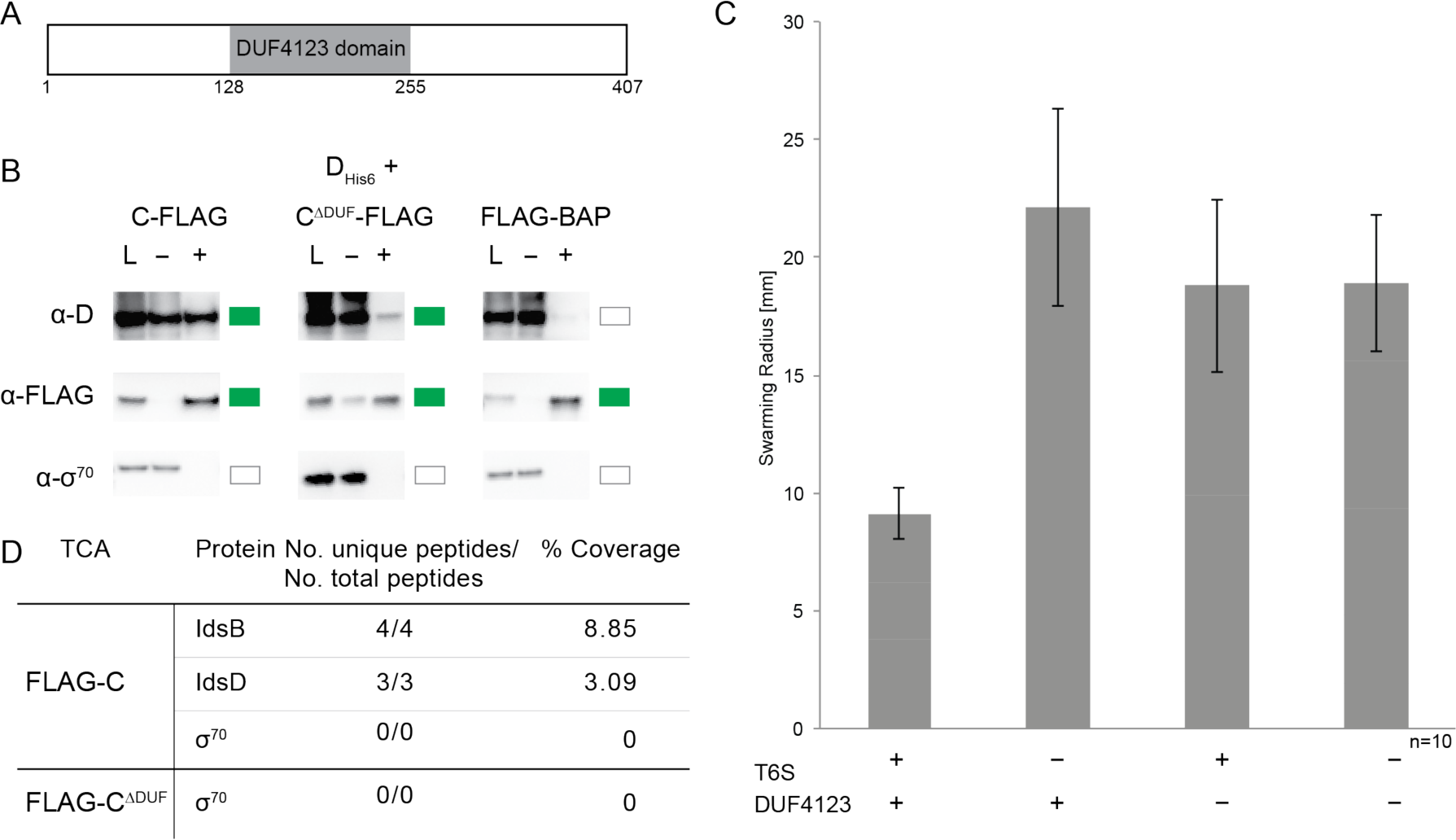
IdsC binds IdsD and is essential for IdsD export. (A) Schematic of IdsC (407 amino acids) drawn to scale; the DUF4123 putative domain spans from amino acids 128 to 255. (B) IdsC-FLAG, IdsC^ΔDUF^-FLAG (IdsC lacking the DUF4123 putative domain), and IdsD-His_6_ (24) were separately expressed in *E. coli* strain BL21(DE3) pLysS from the overexpression vector, pAD100 (55). Lysates were mixed as indicated in the diagram, and anti-FLAG co-immunoprecipitation assays were performed. Soluble (L), non-binding (−) and binding (+) fractions were analyzed via western blot using polyclonal anti-IdsD, monoclonal anti-FLAG, and monoclonal anti-sigma70 antibodies. Green boxes indicate a band in the binding fraction; white boxes indicate no band in the binding fraction. (C) *In vivo* recognition assay for IdsD cell-to-cell transfer in which IdsD export causes a reduced swarm colony radius, while lack of transport results in an increased swarm colony radius (16). DUF4123+ is a *P. mirabilis* strain producing full-length FLAG-IdsC; DUF4123- is a *P. mirabilis* strain producing FLAG-IdsC^ΔDUF^. T6S+ is a *P. mirabilis* strain producing a fully functional T6S system; T6S- is a *P. mirabilis* strain producing a disruption in the TssB homolog that results in disrupted T6S-dependent export (16). Bars indicate migration radii and error bars indicate standard deviations. N=10. (D) Secreted Ids proteins from *P. mirabilis* strains producing either FLAG-IdsC or FLAG-IdsC^ΔDUF^ from a modified pIds_BB_ plasmid (15). The proteins from 75 – 250 kDa of TCA-extracted supernatants from liquid-grown cells were analyzed by LC-MS/MS as previously described (26). The non-secreted sigma-70 protein was used as a control for cell lysis. The presence of three peptides is considered the minimal detectable level.

In the course of our studies, we observed that the mKate-IdsD-associated fluorescence in the IdsC^∆DUF^-containing strain appeared dimmer. As the N-terminal fusion of mKate2 might mask potential effects of the FLAG-IdsC^∆DUF^ mutation, we examined the relative levels of unlabeled IdsD in cell extracts from strains lacking one or more of the Ids proteins and found that unlabeled IdsD was lower or absent in strains completely lacking IdsC or IdsD (Figure 2B). Thus IdsC was not only essential for IdsD subcellular localization into foci, but also for the stable presence of IdsD.

### The IdsC-IdsD interaction is essential for IdsD export

Given these results, we hypothesized that IdsC (Figure 3A) and IdsD likely interact with each other. IdsC and IdsD interactions were examined in *Escherichia coli* strain BL21(DE3) pLysS where no Ids or T6S proteins are otherwise found (52). IdsD fused to a His_6_ epitope tag (IdsD-His_6_) (24) and IdsC fused to a FLAG epitope tag (IdsC-FLAG) were separately produced in this *E. coli* strain. Cell extracts were isolated, mixed, and subjected to anti-FLAG co-immunoprecipitation assays followed by western blot analysis. IdsC-FLAG pulled down IdsD-His_6_ (Figure 3B). The negative control, FLAG-BAP, which is *E. coli* bacterial alkaline phosphatase fused to an N-terminal FLAG epitope, did not pull down IdsD-His_6_ (Figure 3B). These results indicated that IdsC and IdsD bind independently of other Ids and T6S proteins.

As the molecular function of the major domain within IdsC remains unknown, we considered whether the DUF4123 domain (Figure 3A) is essential for IdsC-IdsD interactions. A derivative of IdsC-FLAG without the DUF4123 domain (IdsC^∆DUF^-FLAG) was constructed, produced in *E. coli*, and then subjected to co-immunoprecipitation assays as above. Considerably less IdsD-His_6_ was detected in the pull-down with IdsC^∆DUF^-FLAG than with IdsC-FLAG (Figure 3B). Thus IdsC and IdsD bound each other independently of other Ids and T6S proteins, and the DUF4123 domain was crucial for this IdsC-IdsD interaction.

We reasoned that as IdsC was essential for the subcellular localization of IdsD (Figure 1E) and for IdsD recognition activity (15), then an interaction with IdsC was likely required for IdsD transport itself. This hypothesis was strengthened by evidence that DUF4123-containing proteins are essential for export of T6S substrates in other bacteria (35, 40, 41). To quantify the contribution of IdsC-IdsD binding for export, we assessed *P. mirabilis* strains producing either full-length FLAG-IdsC or FLAG-IdsC^∆DUF^. The resultant strains were subjected to an established *in vivo* recognition assay. In this assay, export of IdsD into neighboring cells that lack IdsE results in a small colony radius; this restricted colony migration is alleviated upon disruption of IdsD export (16). *P. mirabilis* strains producing FLAG-IdsC displayed a small colony radius (Figure 3C), indicating successful export of IdsD (16). By contrast, *P. mirabilis* strains producing FLAG-IdsC^∆DUF^ displayed a large colony radius similar to strains lacking T6S activity (Figure 3C). Abolishing T6S activity in the FLAG-IdsC^∆DUF^ strain displayed no synergistic effects (Figure 3C). As a complimentary approach, we utilized an *in vitro* export assay. The extracellular supernatants of liquid-grown *P. mirabilis* strains producing either FLAG-IdsC or FLAG-IdsC^∆DUF^ were concentrated using trichloroacetic acid precipitations and examined by liquid chromatography tandem mass spectrometry. IdsD is detected in the supernatant of wild-type *P. mirabilis*, but is absent in supernatants of strains lacking T6S function (26, 44). We identified IdsD in the extracellular extract for the strain producing FLAG-IdsC (Figure 3D) but not in that of the strain producing FLAG-IdsC^∆DUF^ (Figure 3D). These results are also consistent with the decreased IdsD protein levels in *P. mirabilis* when full-length IdsC is absent. Thus, we concluded that full-length IdsC was essential for IdsD export.

### IdsC binding to and localization of IdsD is uncoupled from IdsD export

Comparison of the full-length IdsC sequence from the *P. mirabilis* strain used in this study, BB2000, and a strain recognized as non-self, HI4320, showed a uniform length (407 amino acids) with only three single amino acid polymorphisms (99% pairwise identity) (Figure SF6). Comparing the BB2000 IdsC to the HI4320 IdsC, these amino acids are a serine to proline change at position 38 (S38P), a threonine to methionine change at position 121 (T121M), and an arginine to glutamine change at position 186 (R186Q).

We hypothesized that these IdsC residues might be important for binding and export of IdsD, which contains a strain-specific variable region essential for its binding to its self-recognition partner IdsE. We decided to focus on the S38P and R186Q polymorphisms. Lysates from T6S-functional *P. mirabilis* BB2000-derived strains producing either FLAG-IdsC or a mutant variant, FLAG-IdsC^S38P/R186Q^, were subjected to anti-FLAG co-immunoprecipitation followed by western blot analysis for interactions with IdsB (VgrG), IdsD, and IdsE. Both FLAG-IdsC and the FLAG-IdsC^S38P/R186Q^ mutant variant pulled down IdsB and IdsD (Figure 4A). IdsE was largely absent from elutions with FLAG-IdsC; however, there was evidence of IdsE in elutions with the FLAG-IdsC_S38P/R186Q_ mutant (Figure 4A). The FLAG-IdsC and FLAG-IdsC^S38P/R186Q^ constructs were next separately moved into a *P. mirabilis tssB*-disrupted strain that produces mKate-IdsD. Using epifluorescence microscopy, we observed that mKate-IdsD-associated fluorescence was localized into foci in both strains cells (Figure 4B). Therefore, FLAG-IdsC^S38P/R186Q^ binds IdsD and supports IdsD cluster formation.

**Figure 4.**
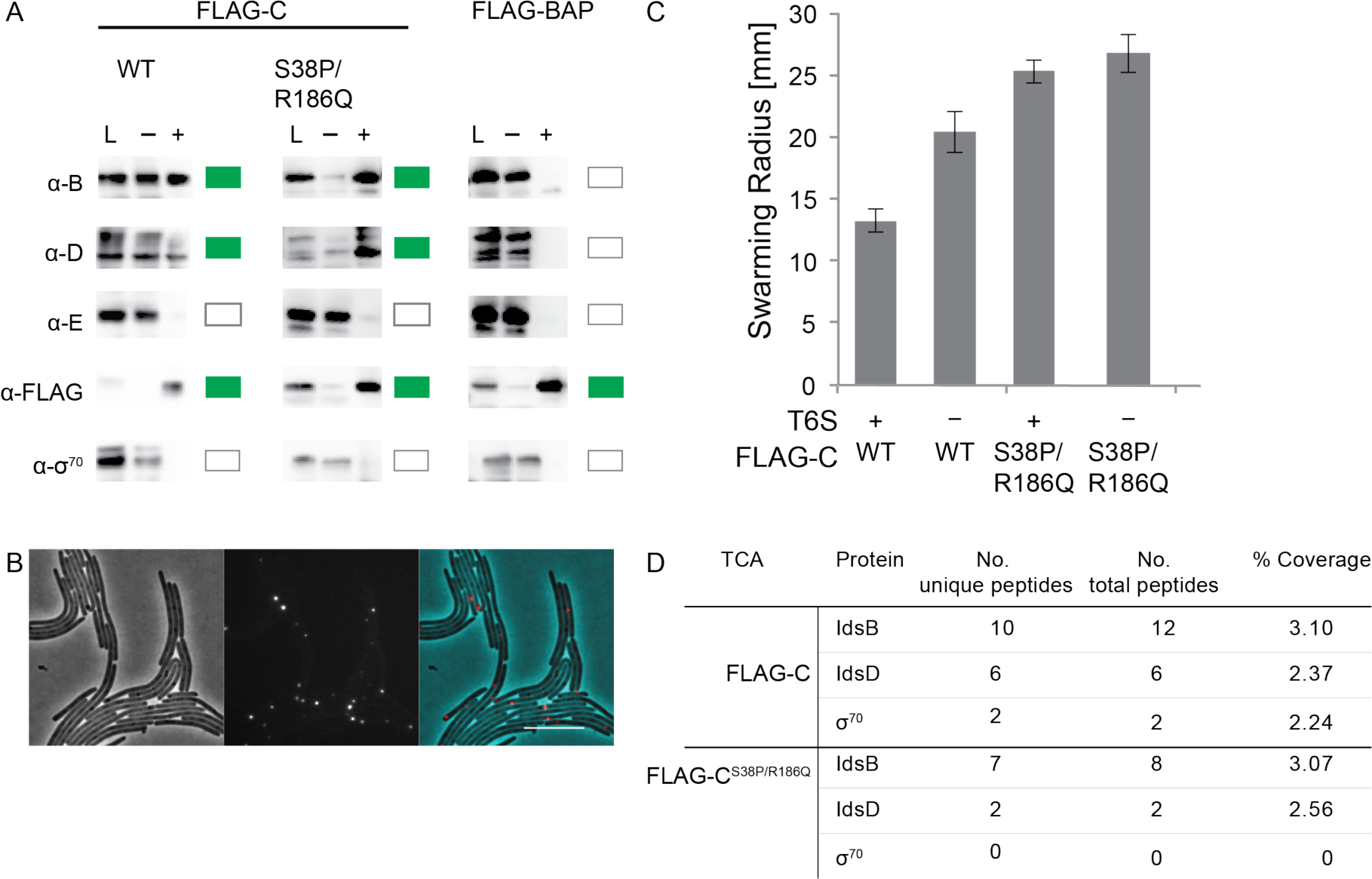
IdsC binding and clustering of IdsD can be uncoupled from IdsD export. (A) FLAG-IdsC or FLAG-IdsC^S38P/R186Q^ were produced in *P. mirabilis*. Lysates were subjected to anti-FLAG batch co-immunoprecipitations. Soluble (L), non-binding (−) and binding (+) fractions were analyzed via western blot using polyclonal anti-IdsB, polyclonal anti-IdsD, monoclonal anti-FLAG, and monoclonal anti-sigma70 antibodies. Green boxes on right indicate a band in the binding fraction; a white box indicates no band in the binding fraction. (B) A *P. mirabilis* strain with disrupted sheath formation producing FLAG-IdsC _S38P/R186Q_ and mKate-IdsD. Left, Phase. Middle, fluorescence in the RFP channel for mKate2. Right, false-colored overlay in which for contrast, mKate-IdsD fluorescence is in red and phase is in cyan. Scale bar is 10 µm. (C) *In vivo* recognition assay for IdsD cell-to-cell transfer in which IdsD export causes a reduced swarm colony radius, while lack of transport results in an increased swarm colony radius (16). T6S+ are *P. mirabilis* strains producing a fully functional T6S system; T6S- are *P. mirabilis* strains producing a disruption in the TssB homolog that results in no T6S-dependent export (16). Error bars represent standard deviations. N = 10. (D). The proteins from 75 – 250 kDa of TCA-extracted supernatants from liquid-grown cells producing FLAG-IdsC or FLAG-IdsC_S38P/R186Q_ from a modified pIds_BB_ plasmid (15) were analyzed by LC-MS/MS as previously described (26). The non-secreted sigma-70 protein was used as a control for cell lysis. The presence of three peptides is considered the minimal detectable level.

In light of these results, we predicted that FLAG-IdsC^S38P/R186Q^ would support IdsD export. We subjected the T6S-functional *P. mirabilis* BB2000-derived strains producing either FLAG-IdsC or FLAG-IdsC^S38P/R186Q^ to the *in vivo* recognition assays as described above. Surprisingly, populations of cells producing FLAG-IdsC^S38P/R186Q^ exhibited an increased colony radius with and without T6S activity, indicating that IdsD was not transferred between neighboring cells (Figure 4C). We used tricholoroacetic acid precipitations of the supernatants from liquid-grown cells to confirm these results. We found that both IdsB and IdsD were detected in the extracellular extract for the strain producing FLAG-IdsC (Figure 4D); however, while IdsB was readily detected, peptides for IdsD were below the detection limit of three peptides for the strain producing FLAG-IdsC^S38P/R186Q^ (Figure 4D). We previously reported that the *in vivo* recognition assay responds to nearly wild-type levels of IdsD transfer and represents a biologically relevant minimum threshold of the transfer necessary for IdsD function (44). These results indicated that IdsC binding of IdsD was uncoupled from IdsD export. Further, T6S export of IdsB was not affected by the two amino acid changes, but the specific export of IdsD was abrogated, strongly supporting that IdsC functions to both bind and sequester IdsD into subcellular clusters and to separately mediate IdsD export via the T6S machinery.

## Discussion

A chaperone-like function for DUF4123-containing proteins encoded next to T6S effectors was recently proposed based on interactions partners and transport efficiencies (40, 41). Our data solidifies this proposed role for DUF4123-containing proteins. We have also meaningfully expanded this proposal by specifically defining a mechanistic model (Figure 5) in which IdsC, a DUF4123-containing protein, binds and stabilizes the T6S substrate, IdsD, independently of transport. We have shown that in addition to the essentiality of IdsC for IdsD transport, IdsC is crucial for the stability and subcellular localization of IdsD.

**Figure 5.**
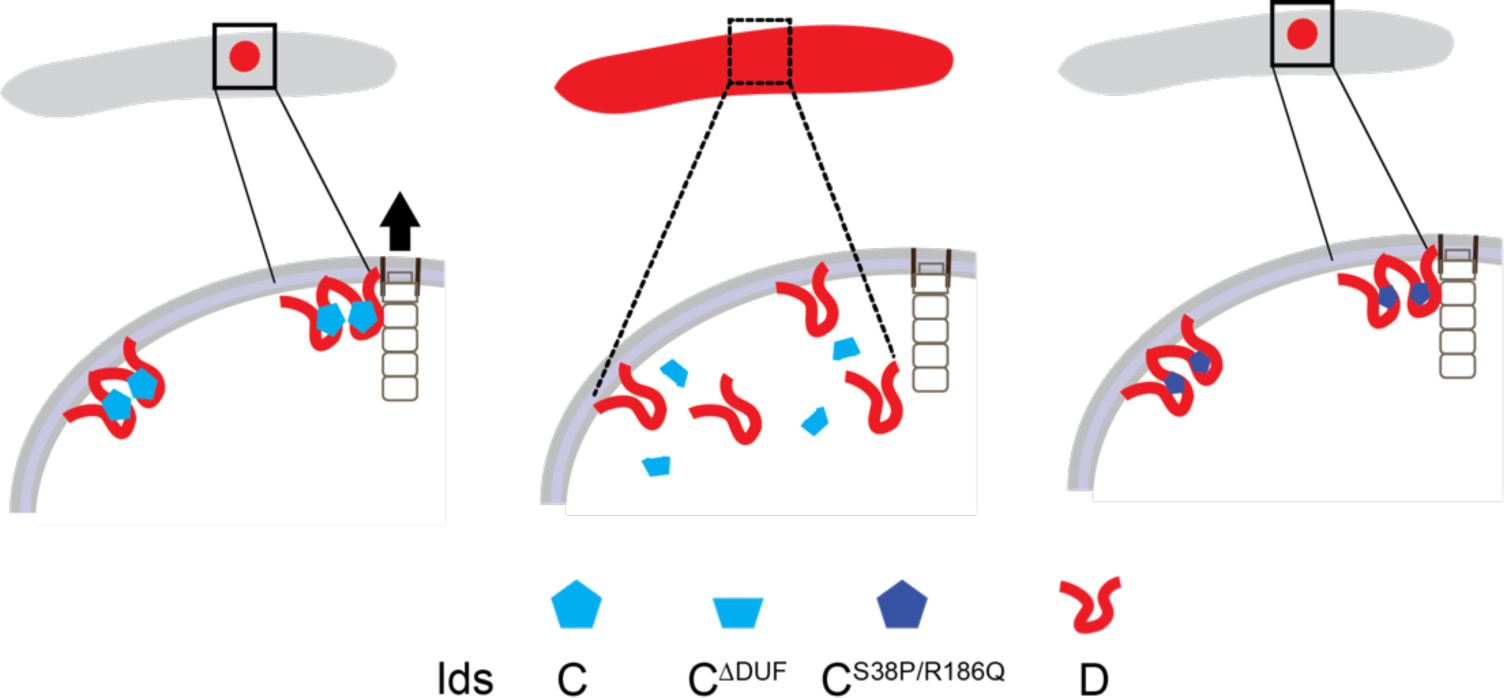
Model for IdsD regulation before delivery into a neighboring cell. We propose that IdsC binds and stabilizes IdsD and that IdsC mediates IdsD transport at the T6S machinery. We posit that aggregation of IdsD by IdsC provides a mechanism to explain the prevention of identity-defining IdsD-IdsE interactions in the donor cell.

The molecular mechanism by which IdsC delivers IdsD for export still remains to be tackled and will likely need to be informed by tertiary structures, none of which are currently available. We hypothesize that within a single cell, IdsD likely binds to IdsB (VgrG) and associates with the T6S membrane-bound complex. We predict that IdsC might protect the transmembrane domains of IdsD from spurious insertion into the donor cell’s inner membrane when IdsD is not docked in the T6S machinery. We further hypothesize that IdsC might actually directly hand IdsD over to IdsB and/or the T6S proteins when the IdsC-IdsD complex is localized at the T6S machinery. In such a case, the amino acid changes in IdsC might alter potential interactions of IdsD with proteins of the core T6S machinery.

Indeed, many organisms face the challenge of inhibiting T6S-exported substrates from acting in the producing cell. The mechanism elucidated here would allow IdsC to prevent IdsD from interacting with its identity partner protein, IdsE, in the donor cell. We propose that both subcellular localization and transport through the T6S machinery function intrinsically to prevent substrate activity within the donor cell and that DUF4123-containing proteins are essential for directing this subcellular regulation. Of note, this proposed mechanism for effector regulation is distinct from previously described models in which an immunity protein is thought to prevent a T6S-exported substrate from acting in the donor cell (30, 33, 42, 43). We surmise that a DUF4123-mediated sequestration might be generally applicable.

This proposed mechanism for IdsC function is reminiscent of chaperone activities in other bacterial transport systems; and yet, several questions remain. Clusters of IdsD are present regardless of the transport machinery, raising the possibility that IdsC might gather IdsD into clusters when IdsD is not actively transported. These IdsD clusters could be for future delivery or alternatively, for sequestration of excess IdsD; such mechanisms require further study. DUF4123-domain proteins are commonly found upstream of T6S substrates even though the sequences for the predicted substrates are quite divergent (40). For the DUF4123-containing proteins studied thus far, the DUF4123-domain proteins appear specific for the T6S substrate encoded immediately downstream (35, 40, 41). Naturally occurring amino acid variations within IdsC alleles did not prevent the binding and subcellular clustering of IdsD before transport, but did impact its export. These results highlight that regulation of T6S substrates before transport might be separate from their loading onto, or transport via, the T6S machinery. Further, DUF4123-containing proteins might be intermediaries between resting and actively transported substrates.

Given that such mechanical aspects are yet unknown, further structural studies dissecting the role of individual amino acid residues within IdsC for binding and export of IdsD are required. More generally, further studies are required to determine whether a specific feature within the DUF4123 domain may be the critical marker for association with a specific T6S substrate and how specificity of substrate binding is obtained.

In conclusion, this research has also provided crucial insights into the communication of self-identity within a population. Prior research left us with the perplexing conclusion that IdsD only acts in recipient cells (15, 16, 24). Here we propose a simple model for the lack of IdsD activity within the donor cell: IdsC appears to couple IdsD sequestration with its localization. We posit that IdsD might exist in two conformations with distinct functions: in a complex including IdsC in the donor cell and in a self-identity complex with IdsE in the recipient cell (16, 24). Such a molecular mechanism (Figure 5) would restrict the communication of self-identity to occur between neighboring cells.

## Author Contributions

M.Z.R, C.C.S, and K.A.G designed experiments; M.Z.R and C.C.S conducted experiments; M.Z.R, C.C.S and K.A.G wrote and edited manuscript.

## Acknowledgements

We thank Lia Cardarelli for contributing experimental materials to this project, as well as members of the Gibbs, Losick, and Gaudet laboratories for thoughtful advice on the project and manuscript. This research was funded by a Howard Hughes Medical Institute Gilliam Fellowship (M.Z.R.) and by Harvard University, the George W. Merck fund, and the David and Lucile Packard Foundation.

## Materials and Methods

### Bacterial strains and media

Strains are described in Table ST1. *E. coli* strains were maintained on LB agar, and *P. mirabilis* strains were maintained on LSW-agar (53). *P. mirabilis* was grown on CM55 Blood Agar Base agar (Oxoid, Basingstoke, England) for swarm assays. For broth cultures, all strains were grown in LB broth under aerobic conditions at 16°C, 30°C, or 37°C. Antibiotics were used at the following concentrations: 100 microgram/milliliter (µg/ml) carbenicillin, 15 µg/ml tetracycline, 35 µg/ml kanamycin, 50 µg/ml chloramphenicol, and 25 µg/ml streptomycin.

### Strain construction

All chromosomal mutations in BB2000 and *∆ids* were made as described in (16) using pKNG101-derived suicide vectors (54). The *tssK*_*null*_ strain introduced a *BB2000_0814*_*G1329T*_ mutation using plasmid pCS33a. The *tssK*_*partial*_ strain introduced a *BB2000_0814*_*G1145A*_ mutation using plasmid pCS33b. *BB2000_0814* (GenBank accession no. AGS59310.1) is a gene encoding a TssK homolog (T6SS_VasE PFAM family PF05936). Mutations in the *∆ids* background were confirmed via whole genome sequencing by the Bauer Core Facility at Harvard University using the protocol described in (16). Mutations into the BB2000 background were confirmed by Polymerase Chain Reaction (PCR) amplification of *BB2000_0814* (primers: 5′- CTCTCCGGCAATAATACGTAG-3′ and 5′- CAGACCCACTACAGGCTTTAG-3′) followed by Sanger sequencing performed by GENEWIZ, Inc. (South Plainfield, NJ).

### Plasmid construction

Plasmids and construction details are described in Table SF2. Restriction digest and subsequent ligation of gBlock gene fragments (Integrated DNA Technologies, Coralville, IA) were used for cloning of both pIds_BB_-derived (15) and pAD_100_-derived (55) vectors. gBlock gene fragments were first subcloned into TOPO pCR2.1 vector using the TOPO TA-Cloning Kit (Thermo Fisher Scientific, Waltham, MA). All methods were performed according to manufacturers’ instructions. The FLAG epitope is DYKDDDDK and was introduced using Quikchange reaction protocols (Agilent Technologies, Santa Clara, CA). The pIds_BB_-derived plasmids were constructed via restriction digest using listed restriction enzymes (New England BioLabs, Ipswich, MA). Ligations were resolved in OneShot Omnimax2 T1R competent cells (Thermo Fisher Scientific, Waltham, MA). Resultant plasmids were confirmed by Sanger sequencing (Genewiz, Inc., South Plainfield, NJ) and correct resultant plasmids were then transformed into *P. mirabilis* as described (15). All *P. mirabilis* constructs were tested for functionality using a standard boundary formation assay (56). Ligations for plasmids derived from pAD_100_ (55) were resolved in XL10-Gold Ultracompetent cells (Agilent Technologies, Santa Clara, CA). Correct resultant plasmids were transformed into BL21(DE3) pLysS (Thermo Fisher Scientific, Waltham, MA). A flexible linker (GSAGSAAGSGEF, (57)) was used to separate the mKate2 fluorophores from IdsD. All vectors were confirmed by sequencing with site-specific primers using the services of GENEWIZ, Inc. (South Plainfield, NJ).

### α-FLAG immunoprecipitation assays

Assays were performed and analyzed as previously described (24). Modifications to those protocols are as follows. *P. mirabilis* cells were harvested from swarm-permissive media after incubation at 37°C for 16–20 hours. Control lysate (containing pIds_BB_ with no FLAG-tagged protein) was supplemented with 2 µg of purified FLAG-BAP protein (Sigma-Aldrich, St. Louis, MO). *E. coli* BL21 (DE3) pLysS cells (Thermo Fisher Scientific, Waltham, MA) were grown in 25 mL of LB supplemented with carbenicillin under shaking conditions at 30°C until optical density at 600 nm (OD_600_) was between 0.6 and 1. Cultures were cooled on ice, induced with 1 mM IPTG, and incubated overnight shaking at 16°C. Cells were harvested by centrifugation and stored at −80°C. Lysates were mixed to a total volume of 1 ml of which 900 µl was applied to 40 µL pre-equilibrated α-FLAG M2 antibody resin (Sigma-Aldrich, St. Louis, MO; Biotools, Houston, TX). *P. mirabilis* and *E. coli* extracts were obtained separately as described above. Lysates were mixed to a total volume of 1 ml of which 900 µl was applied to 40 µL pre-equilibrated α-FLAG M2 antibody resin (Sigma-Aldrich, St. Louis, MO; Biotools, Houston, TX).

### Antibody production

Antibodies specific to IdsB amino acids 713–723 (CRAKAMKKGTA), IdsD amino acids 4–18 (EVNEKYLTPQERKAR) (24), and IdsE amino acids 298–312 (EQILAKLDQEKEHHA) (24) were raised in rabbits using standard peptide protocols (Covance, Dedham, MA).

### SDS-PAGE and western blots

Assays were performed and analyzed as previously described (24). Polyclonal primary antibody dilutions were rabbit α-IdsB (1:5000), rabbit α-IdsD (1:2000), and rabbit α-IdsE (1:2000). Monoclonal primary antibody dilutions were rabbit α-FLAG (1:4000, Sigma-Aldrich, St. Louis, MO) and mouse α-σ^70^ (1:1000, Thermo Fisher Scientific, Waltham, MA or BioLegend, San Diego, CA). Secondary antibodies were goat α-rabbit or goat α-mouse antibodies conjugated to HRP (polyclonal, 1:5000, SeraCare Life Sciences, Milford, MA).

### Trichloroacetic acid precipitations (TCA) and liquid-chromatography tandem mass spectrometry analysis (LC-MS/MS)

All trichloroacetic acid precipitations were performed as previously described (26). Binding fractions from α-FLAG immunoprecipitations or supernatant fractions from TCA were separated by gel electrophoresis using 12% Tris-Tricine polyacrylamide gels and stained with Coomassie blue as previously described (16, 26). Supernatant fractions from TCA were cut into two bands at 75–150 kDa and 10–25 kDa. LC-MS/MS was performed by the Taplin Mass Spectrometry Facility (Harvard Medical School, Boston, MA). Bioinformatics analysis of Ids and T6S protein hits was done using Pfam 31.0 (58).

### Colony expansion

Colony expansion assays were conducted as previously described (16). Modifications to those protocols are as follows. Swarming-permissive plates supplemented with kanamycin were inoculated with 1 µl of overnight cultures normalized by OD_600_. Plates were incubated at 37° for 17 hours and swarming radii recorded.

### Microscopy

Strains were grown overnight shaking in LB supplemented with kanamycin at 37°. 1.0mm-thick agar pads of swarming permissive media supplemented with kanamycin were inoculated with 2 µl of overnight cultures and incubated in a humidified chamber at 37° for 4 to 6 hours. Images were acquired either with a Leica DM5500B (Leica Microsystems, Buffalo Grove, IL) with a CoolSnap HQ^2^ cooled CCD camera (Photometrics, Tucson, AZ) or an Olympus BX61 (Olympus Corporation, Waltham, MA) with a Hamamatsu C10600-10B CCD camera (Hamamtsu Phototonics K.K., Boston, MA). MetaMorph version 7.8.0.0 (Molecular Devices, Sunnyvale, CA) was used for image acquisition. Figures were made in Fiji (59, 60) and Adobe Illustrator (Adobe Systems, San Jose, CA).

